# Open-eyed meditation suppresses functional connectivity in EEG across a broad frequency range

**DOI:** 10.64898/2026.05.13.724915

**Authors:** Vinod Jiani, Ankan Biswas, Supratim Ray

**Author notes:** **Conflict of interest** The authors declare no competing financial interests. **Acknowledgments** We thank Srishty Aggarwal and Dr. Kanishka Sharma for their assistance in data collection. We acknowledge the use of LLMs for grammar and literature review. **Author contributions** Conceptualization: VJ, AB, SR; Methodology: VJ, AB, SR; Experimentation: AB; Supervision: SR; Writing: VJ, AB, SR.

## Abstract

Functional connectivity (FC) is a statistical measure that reflects the degree of phase consistency between two signals and provides insights about potential interactions between two brain regions. Previous studies have reported conflicting results on the effect of meditation on FC, with some showing enhancement while others reporting suppression of FC. However, even though meditation increases power over a broad frequency range between 15-200 Hz and beyond, most FC studies have reported changes over fixed and narrow frequency bands below 50 Hz. Further, meditation-induced changes in power spectral density (PSD) and FC have never been compared with changes with other factors such as age, gender and stimulus. We recorded electroencephalogram (EEG) from open-eyed meditators (N=35) and their gender-and age-matched controls (N=36) and found that meditation was associated with a‘state’ decrease in FC across a broad frequency range (15-200 Hz), while PSD showed both ‘trait’ and ‘state’ enhancement. Furthermore, visual gratings, which are known to enhance narrow-band gamma power, led to reduced gamma FC in both meditators and controls. We also compared the effect of aging and gender on a different dataset of healthy middle-aged (N=78) and elderly (N=89) participants and found differences in distinct frequency bands that were limited to a narrow range. We also found that often-used average referencing heavily distorted the FC and gave uninterpretable results. Overall, our results suggest distinct neural mechanisms underlying healthy aging, vision, and meditation and further recommend caution while using average referencing to study phase-based metrics.

**Significance statement:** Meditation research has reported inconsistent effects on functional connectivity (FC), partly because most studies examined only narrow low-frequency bands despite meditation altering brain activity across a much broader frequency band. This study demonstrates that meditation produces a broadband state reduction in FC across 15–200 Hz, while simultaneously enhancing power. In contrast, healthy aging, gender, and visual stimulation showed frequency-specific effects confined to alpha (8-12 Hz) and high-beta (20-36 Hz) bands, highlighting meditation’s unique large-scale neural signature. The study also shows that average referencing can severely distort phase-based FC estimates, leading to misleading interpretations. These findings clarify conflicting literature, distinguish meditation from other neural modulators, and provide important methodological guidance for EEG connectivity research.

## Introduction

Meditation encompasses a wide spectrum of contemplative practices known to improve attention, mitigate stress, enhance emotional regulation and overall mental health, and help in the treatment of psychiatric disorders (Ivanovski and Malhi, 2007; Tang et al., 2007; Sumantry and Stewart, 2021). Many studies have shown that the power spectral density (PSD) of the electroencephalogram (EEG) of meditators is different from age and gender matched controls, even while the subjects are not meditating (i.e., a “trait” effect; (Cahn and Polich, 2006; Lomas et al., 2015)). In addition, PSD also changes during meditation (a “state” effect), with a prominent increase in power in a broad frequency range above ∼15 Hz to up to 200 Hz and beyond (Ferrarelli et al., 2013; Braboszcz et al., 2017; Biswas et al., 2026).

There has also been a growing interest in investigating coupling between different brain areas during meditation using functional connectivity (FC) – a statistical measure reflecting the extent of phase consistency between two signals (Shen et al., 2020). However, most studies to date have investigated changes in FC in fixed frequency bands below 50 Hz, even though the corresponding changes in power are broadband (Biswas et al., 2026). Further, different meditation practices have been associated with distinct alterations in EEG FC (Cahn and Polich, 2006; Ivanovski and Malhi, 2007). One study found reduced FC between cortical sources in meditators of five traditions in the frequency range between 1.5-44 Hz (Lehmann et al., 2012), whereas another study reported enhanced long-distance synchrony in the gamma band (25-42 Hz) in meditators practicing loving-kindness (Lutz et al., 2004). Another study found lower gamma FC (25-45 Hz) for meditators compared to controls during the resting state in frontal and posterior regions (Berkovich-Ohana et al., 2014). However, no study to date has shown FC across the spectrum up to 200 Hz. Further, previous studies have reported FC changes with closed-eye meditation. Also, meditation practiced with open eyes remains understudied.

Several other factors, such as stimulus presentation, age, and gender, also influence the PSD and FC in EEG. While one study reported reduction in FC in alpha and gamma bands with aging (Kumar and Ray, 2023), others have reported mixed results (Sifis et al., 2009; Perinelli et al., 2022). One study reported gender differences in FC in sleep, with females showing higher high-sigma FC (14-16 Hz) but lower alpha (8-13 Hz) and beta (20-30 Hz) FC than males (Ujma et al., 2019). Importantly, no study has compared the effect of meditation on FC with that of healthy aging, gender, and visual stimulus. Further, all previous studies showed FC calculated in pre-specified narrow bands, so how different conditions affect the full FC spectrum remains unknown.

FC is strongly dependent on the referencing scheme (Essl and Rappelsberger, 1998; Shirhatti et al., 2016; Ríos-Herrera et al., 2019), and several studies suggest that average referencing of the signals is preferred over unipolar referencing since it reduces bias from a single physical reference electrode (Srinivasan et al., 2007; Lehmann et al., 2012; Ríos-Herrera et al., 2019) and improves comparability across studies (Huang et al., 2017). However, we have previously shown that average referencing heavily distorts the phase relationship in local field potential data (Shirhatti et al., 2016). Whether average referencing affects EEG FC as well is unknown.

To address these gaps, we studied FC on EEG data from advanced practitioners of open-eyed Rajyoga meditation (RYM) and their age-and gender-matched controls while they were presented with visual stimuli before, during, and after meditation. Using another dataset in which stimulus-induced gamma was studied on a large cohort of 89 middle-aged and 78 elderly participants, we also tested how aging and gender affect the full FC spectrum. To study the effect of referencing, we repeated the same analyses on common-average-referenced signals. We also analyzed lagged coherence and imaginary coherence to address the problem of zero-phase lag that often arises from volume conduction (Nolte et al., 2004; Pascual-Marqui, 2007).

## Materials and Methods

### Meditation Dataset

We collected EEG data from 78 participants, primarily from the local community in Bengaluru, aged 20-65 years. The cohort included 38 experienced meditators (19 female) with at least 5 years of regular meditative practice, and 40 control participants (17 female) with no prior meditation experience. All individuals completed a subjective screening questionnaire that assessed chronic or current illnesses and medication use to ensure participants were in good health. For female participants, we also noted their menstrual cycle status at the time of data collection. Participants also confirmed via self-report that they had not consumed alcohol, tobacco, or any recreational drugs within the two weeks preceding the experiment.

The meditators were trained in a tradition called Raja Yoga Meditation by the Brahma Kumaris (BK) organization. It is practiced with open eyes. The meditators were given a questionnaire designed by the parent organization (BK) to assess their total hours of meditation experience and other factors related to lifestyle and dedication to the meditation practice. A top-down coordination from the Mt. Abu Headquarters of BK to their local centers in Bengaluru was done to facilitate data collection. The participants who were recommended by the local center administrators were invited to the experiment.

Control participants of the same gender and age (±2 years) were invited locally through posters, broadcast emails, and word-of-mouth. We tried to schedule the experiment for female participants in such a way that the phase of the menstrual cycle of a meditator was close to that of their control (or vice versa), since alpha and gamma oscillations are modulated by menstrual phase (Sumner et al., 2018). Informed consent was obtained from all participants, and they were provided with monetary compensation for their time. All procedures were approved by Institutional Human Ethics Committee of Indian Institute of Science, Bengaluru.

### Participant selection

Participant selection is described in detail in our previous study (Biswas et al., 2026) and briefly discussed here. We first used an automated pipeline to identify bad electrodes and trials for each participant. We rejected 5 participants (1 meditator and 4 controls) who had more than 24 (out of 64) bad electrodes (See Artifact Rejection subsection below for more details). Hence, we were left with usable data from 71 participants: 35 meditators (17 female) and 36 controls (16 female). We paired each meditator with a control of the same sex in a way that minimized the difference – in order of preference – in age, education level, menstrual phase (for females), and experiment date. Finally, we could find appropriate controls for 30 meditators (14 female). The demographics of the participants are described in detail in our previous study (Biswas et al., 2026). We have interchangeably used the words ‘sex’ and ‘gender’ in this study, denoting the biological sex of the participants.

### Tata Longitudinal Study of Aging (TLSA) dataset

As the number of participants in the meditation dataset was insufficient to study the effect of aging and gender on FC, we used the TLSA dataset described in detail in our previous study (Murty et al., 2020, 2021), in which participants performed a visual fixation task, where full-screen grating stimuli were presented for 800 ms with an inter-stimulus interval of 700 ms after a brief fixation of 1000 ms in each trial. The rest of the Methods describes the Meditation experiment, although the data acquisition system and analyses pipelines are very similar across both studies (see (Murty et al., 2020, 2021) and (Kumar and Ray, 2023) for more details)).

### EEG data collection

EEG data were acquired using 64 active electrodes (actiCAP) and a BrainAmp DC system (Brain Products GmbH), with electrodes positioned according to the international 10–10 system and referenced online to FCz (we refer to this as the ‘unipolar’ reference scheme). We filtered the signals online between 0.016 Hz (first order) and 250 Hz (fifth-order Butterworth), sampled at 1000 Hz, and digitized at 16-bit resolution (0.1 μV/bit). Participants sat in a dark room facing a gamma-corrected LCD monitor (BenQ XL2411; 20.92 × 11.77 inches, 1280 × 720 pixels, 100 Hz refresh rate) positioned ∼58 cm away, subtending 49° × 29° of visual angle for full-screen gratings. Eye position was recorded at 1000 Hz with an EyeLink Portable Duo head-free eye tracker (SR Research Ltd).

### Experimental procedure

The experiment procedure is described in detail in our previous study (see Figure 1 of (Biswas et al., 2026)) and briefly mentioned here. It comprised of eight protocols. It began with eyes-open fixation at a centre dot on the screen (EO1), then an eyes-closed state (EC1), followed by a grating protocol (G1) in which full-screen gratings were shown with passive fixation for 1.25 s with equal interstimulus intervals. Stimuli were continuous, and (if needed) participants blinked during interstimulus periods. Each of EO1, EC1, and G1 lasted ∼5 minutes. This was followed by open-eye meditation with gaze maintained on the screen for ∼15 min (M1). The first three protocols were then repeated (G2, EO2, EC2), and the final protocol (M2) involved meditation with open eyes while gamma-inducing gratings (as in G1/G2) were presented for ∼15 min. Gratings were achromatic, with spatial frequencies of 2 or 4 cycles per degree (cpd) and orientations of 0°, 45°, 90°, 135° (8 combinations). Even for EO1, EC1, EO2, EC2, and stimulus-free meditation (M1), we segmented the data into 2.5-second “trials” (1.25 s baseline + 1.25 s stimulus) to match trial structure; for non-stimulus protocols, PSDs, and FCs from both epochs were averaged. All protocols had 120 trials except M1 and M2 (360 trials each).

**Figure 1:**
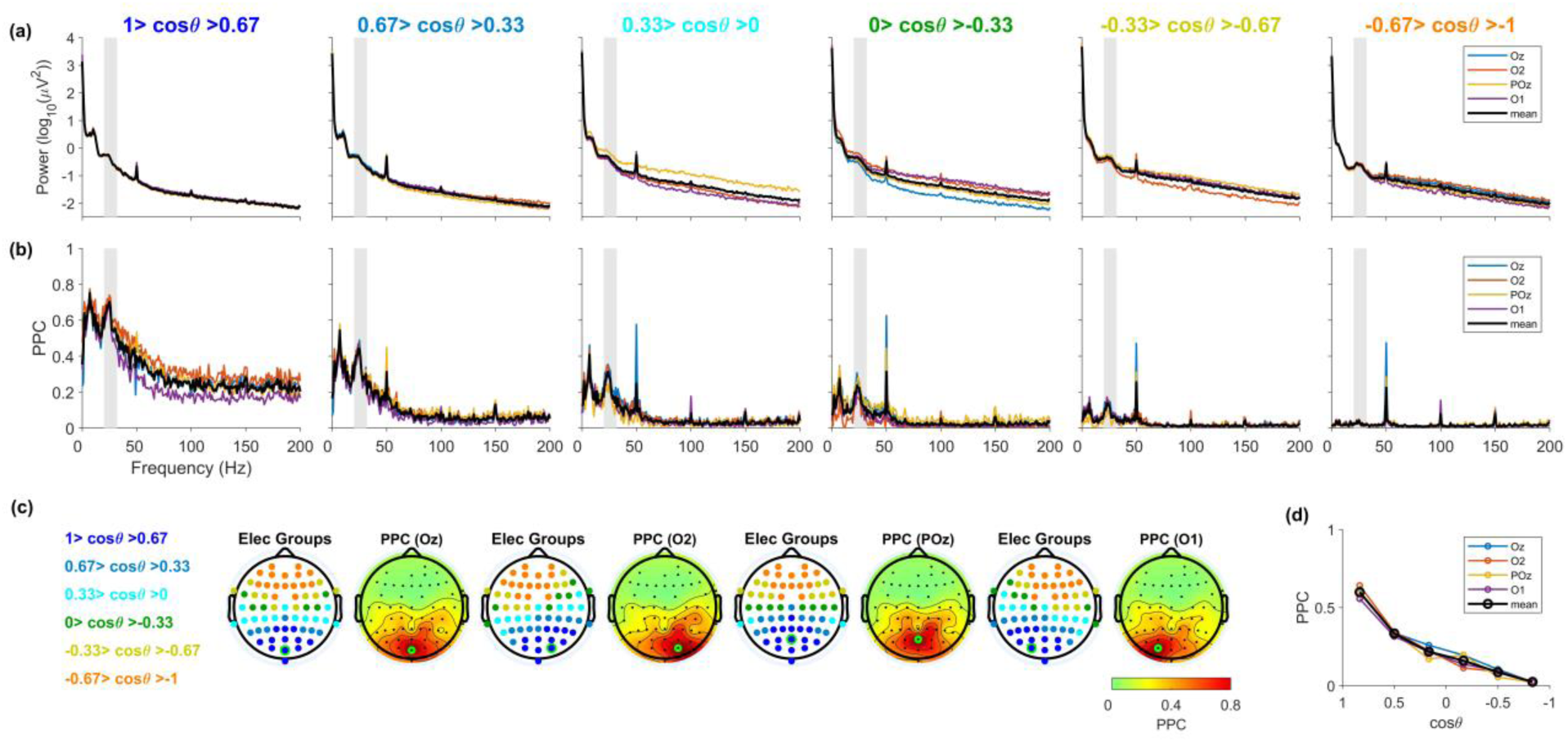
Power and functional connectivity (FC) for a representative participant during EO1: **(a)** The power spectral density (PSD) of different electrode groups defined based on their distance from a ‘seed’ electrode selected from the occipital cluster (Oz, O2, POz, O1) for a sample participant. The corresponding four PSDs are plotted in respective colors, and the average of the four is plotted in bold black color, as indicated in the legend. **(b)** The corresponding FC measured using pairwise phase consistency (PPC) with respect to the four seed electrodes is plotted in the respective colors. The average FC is plotted in bold black color as indicated in the legend. **(c)** The scalp layouts of electrode groups (left) and scalp maps of FC (right) for the four occipital seed electrodes (highlighted in green). The scalp layouts show the six electrode groups defined based on the interelectrode distance from the corresponding seed electrode, highlighted in green (for details, refer to‘Visualizing average FC’ under the’Methods’ section). The electrode groups are color-coded in terms of the cosine of their angular distance (cosθ) from the seed electrodes (mentioned on the extreme left). FC is averaged over the frequency window (20-32 Hz), which is shown by the vertical shaded gray region in (a) and (b). **(d)** FC plotted against the inter-electrode distance relative to the four seed electrodes in respective colors. The FC value for each electrode group is obtained by averaging the PPCs in (b) over the frequency window of interest, 20-32 Hz.

### Meditation technique

#### Meditators

Meditators practiced open-eye Raja Yoga meditation (‘seed-stage’ meditation) of the BK tradition, which involves contemplation and directed thought to attain experiential states for self-development (Nair et al., 2017). The practice centres on shifting awareness from the external world to “the soul and its peaceful nature” and then connecting to the “Supreme Soul.” Practitioners begin with bodily awareness, then focus on or behind the forehead, visualizing themselves as a point of light – “soul consciousness,” a core aspect of BK Raja Yoga meditation (Nair et al., 2017; Sharma et al., 2023). In the final stage, they connect with the Supreme Soul, reflecting on and affirming seed-like qualities – peace, power, bliss, love, purity, and wisdom – imagining these qualities being received and permeating the soul. This induces feelings of completeness, detachment from the physical world, silence, and subtlety. The seed stage represents awakening inherent spiritual qualities, reinforced by affirmations such as “I am a peaceful soul.” With practice, meditators can quickly enter and sustain this state.

#### Controls

To familiarize controls with relaxation practice, they received male-voiced audio instructions in English and Hindi and were asked to practice at least once before the EEG recording. The practice began with body scanning, directing attention from the toes upward to the head, coordinated with breathing, with the instruction that *“every incoming breath is energizing your body, and every outgoing breath is relaxing it”.* Participants then focused on the center of the forehead, visualizing a point of light. This entire sequence is similar to the initiation procedure of BK meditation. During the EEG recording, they performed the same practice without audio guidance to avoid potential auditory stimulation of the brain.

### Artifact rejection

Artifacts were removed following the criteria adapted from (Murty and Ray, 2022). Electrodes with impedance >25 kΩ were rejected; the final set had a mean ± SD impedance of 6.19 ± 4.80 kΩ. For each electrode, trials were flagged as outliers if RMS exceeded 1.5–35 μV or if frequency-domain deviation exceeded 6 SD from the mean. Electrodes with >30% outlier trials were discarded. Trials classified as bad in occipital electrodes or >10% of other electrodes were removed, yielding a common set of bad trials per protocol and participant. This resulted in rejection rates (mean ± SD) of 17.2 ± 6.9%, 19.2 ± 7.0%, 16.8 ± 5.5%, 23.8 ± 7.4%, 17.7 ± 6.5%, 18.5 ± 7.9%, 19.8 ± 8.2%, and 23.1 ± 8.0% for EO1, EC1, G1, M1, EO2, EC2, G2, and M2, respectively. Electrodes with power spectral slopes <0 in the frequency window 56-84 Hz were rejected. Fp1, FC2 to FC6, FT7, TP8, and C2 were excluded from all analyses as they were bad in >35% of participants in either group. Participants with >35% bad electrodes were excluded (5 total; 1 meditator, 4 controls).

#### Eye-artifacts

Eye blinks or gaze shifts beyond ±2.5° from fixation between −1 and 1.25 s of stimulus onset were marked as fixation breaks and removed, leading to rejection rates of 31.9 ± 21.1%, 22.2 ± 19.9%, 39.0 ± 27.3%, 25.7 ± 21.1%, 33.0 ± 23.1%, and 26.2 ± 20.0% for EO1, G1, M1, EO2, G2, and M2, respectively. These higher rates compared to (Murty et al., 2020) are due to the longer analysis window and continuous stimulus presentation. Analyses repeated without removing‘bad-eye-trials’ yielded similar results.

Participants with fewer than 30 good trials and fewer than 3 usable electrodes in each group, for both stimulus and baseline epochs, were excluded. This led to a few additional rejections, leaving 29-30 pairs for most protocols, except G2 (28 pairs). Cross-protocol comparisons (e.g., M1 vs EO1 and M2 vs G2) were restricted to pairs meeting criteria for both protocols, yielding 22–30 matched pairs per comparison.

### EEG data analysis

All analyses were done on the original signals (i.e., using unipolar referencing). However, since previous studies have often used average referencing (Lehmann et al., 2012; Ríos-Herrera et al., 2019), we also performed the same analyses on the data obtained with average referencing, which yielded strikingly different results that are discussed later. We used the Fieldtrip Toolbox (Oostenveld et al., 2011) to generate average-referenced data.

Since functional connectivity is always calculated between two electrodes, i.e., a‘seed’ (reference) electrode and any other electrode, we mainly used occipital seed electrodes (Oz, POz, O2, and O1), as done previously (Kumar and Ray, 2023). We also tried other seed electrodes, such as frontal (Fz, AFz, AF3, AF4, F1, and F2), and got comparable results (Figure S1). We tried many other seed groups, and most of them (such as left temporal, right temporal, left parietal, and right parietal group) showed a significant difference in the FC of meditators and controls, suggesting that the broadband suppression in FC is a widespread phenomenon not limited to the occipital seed electrodes.

We used custom codes written in MATLAB (MathWorks. Inc; RRID: SCR_001622) to carry out all the data analyses. We generated the scalp maps using *topoplot* function of the EEGLAB toolbox (Delorme and Makeig, 2004), RRID: SCR_007292) with standard ActiCap 64 unipolar montage of the channels. Power spectral density was obtained using the Chronux Toolbox (Bokil et al., 2010, RRID: SCR_005547) for individual trials and then averaged over trials for a given electrode. For G1, G2, and M2, the baseline period between-1 and 0 seconds of stimulus onset was considered for analysis, while the stimulus period was taken between 0.25 and 1.25 seconds to avoid stimulus-onset-related transients, giving us 1 Hz of frequency resolution for the PSDs. For uniformity, for other non-stimulus protocols (EO1, EC1, M1, EO2, and EC2), we inserted imaginary stimulus markers at 2.5-second intervals, considered the baseline and (imaginary) stimulus periods as before, and then averaged the PSDs across stimulus and baseline.

The FC measures were calculated using the Fieldtrip Toolbox (Oostenveld et al., 2011), RRID: SCR_004849). It was calculated within the same stimulus period to minimize the influence of stimulus-onset transients. Since FC metrics may depend on the number of trials (Vinck et al., 2010; Miskovic and Keil, 2015), we matched the number of trials across protocols before computing FC metrics. Also, we primarily used the pairwise phase consistency (PPC) (Vinck et al., 2010) as our FC metric, which is free from finite-sample bias (see (Vinck et al., 2010) for details). PPC is calculated as follows.

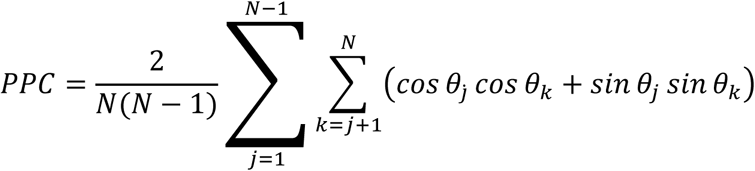

where N is the number of trials. 𝜃_𝑗_ 𝑎𝑛𝑑 𝜃_𝑘_ are relative phases from two signals (Vinck et al., 2010).

Phase differences and phase locking value were estimated using the multi-taper method (Thomson, 1982), in the Chronux Toolbox (Bokil et al., 2010). To obtain maximum frequency resolution, we used a single taper. If the Fourier coefficients of two signals are expressed as 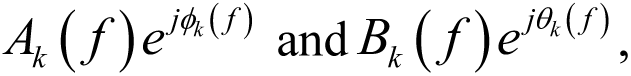 (where f is the frequency and k is the trial number, which varies between 1 and N), phase coherence or phase locking value is calculated by (Lachaux et al., 1999)

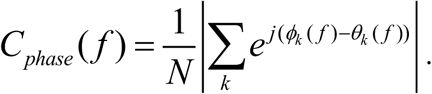

### Visualizing average FC

Following previous literature (Kayser and Tenke, 2015), we computed the inter-electrode distance (IED) in the elevation and azimuth angular space. As a measure of IED, we calculated *cos θ,* where *θ* is the angular separation between the seed electrode and the electrode under consideration. To study how FC varies with IED from the seed electrode to other electrodes, we averaged FC between electrode pairs within defined IED intervals – six intervals covering 180 degrees, each with an angular spread of ∼30 degrees. In the cosine IED space, it is equivalent to six intervals covering [-1, 1], each with a spread (bin width) of 0.33, as shown in Figure 1 (c).

## Statistical analysis

As our data was paired across participants in the two groups, we compared the means of the PSDs and PPCs between meditators and controls using a paired t-test. We used an unpaired t-test for the same comparisons between elderly and middle-aged subjects, and males and females.

## Data and code availability

We performed spectral analyses using Chronux toolbox (version 2.10), available at http://chronux.org. FC measures were calculated using the Fieldtrip Toolbox, available at https://www.fieldtriptoolbox.org/. The codes to see the analyzed power and connectivity data are available on GitHub at github.com/supratimray/ProjectDhyaanBK1Programs.

## Results

We collected EEG data from 78 participants (38 long-term meditators with an average of ∼10,000 hours of meditation practice and 40 controls; see our previous study (Biswas et al., 2026) and Methods for more details. After removing participants with more than 24 (out of 64) bad electrodes, we were left with usable data from 71 participants (35 meditators and 36 controls). We paired each meditator with a control participant of an age within 2 years and obtained 30 matched pairs. Participants with fewer than 30 good trials and two good electrodes per electrode group in each protocol were rejected, so the number of matched pairs of meditators and controls varied between 28 and 30 across analyses. The participant and experiment details are elaborately mentioned in our previous study (Biswas et al., 2026).

EEG data were collected for 8 sessions, named EO1, EC1, and G1 (5 minutes each); M1 (15 minutes); G2, EO2, and EC2 (5 minutes each) and M2 (15 minutes), as described in Fig.1 of our previous study (Biswas et al., 2026). Thus, participants performed open-eyed meditation twice – once without any stimulus (protocol M1) and later while grating stimuli were shown (M2). Eye position was measured using a head-free eye tracker (Eyelink Duo; SR-Research). A fixation spot was shown for all eyes-open segments (EO1, EO2, G1, G2, M1, M2), and epochs when the eye position deviated by more than 2.5 degrees from it were removed later.

### FC analysis with occipital seed electrodes shows distinct peaks in the alpha and beta bands in the resting state

Figure 1 shows the FC analysis pipeline for a sample participant during EO1. We considered four seed electrodes in the occipital region, namely Oz, O2, POz, and O1, as shown in the scalp layouts in Figure 1c. Next, we divided all electrodes into six groups based on the distance from the seed electrode (Figure 1c). We computed the pairwise phase consistency (PPC) – an unbiased estimator of the square of the phase locking value (Vinck et al., 2010) – for each electrode with each seed electrode. We also computed coherence and phase locking value and obtained similar results (not shown). However, the weighted phase leg index (wPLI), lagged coherence, and the imaginary coherence (iCoh) yielded inconsistent results, as discussed later. Figure 1a shows the average power spectral density (PSD) of the electrodes for the six electrode groups. The curves in different colors show the PSDs of the electrode groups, defined relative to the four seed electrodes (Figure 1a). The peaks at 50, 100, and 150 Hz correspond to line noise and harmonics. Figure 1b shows the corresponding FC for the same six electrode groups, averaged over the electrodes within each group.

As expected, we found that FC decreased monotonically with the inter-electrode distance, quantified by cosθ, where θ is the angular separation between the seed electrode and the electrode under consideration (Figure 1b). Interestingly, we found a prominent‘bump’ in the FC in the beta frequency range between 20-32 Hz (highlighted in gray shadow in Figure 1a and b), in addition to the alpha bump at 8-12 Hz. To analyze the spatial spread of FC in the former frequency window, we plotted scalp maps of FC relative to each seed electrode (Figure 1c). We also plotted the FC against inter-electrode distance (cosθ) (Figure 1d). Since the FC showed a similar distribution relative to each of the four seed electrodes (Figure 1b, c, d), we averaged the FC over this local cluster of occipital seed electrodes in all subsequent analyses (shown as black traces here in all plots).

### During non-meditative states, FC is similar between meditators and controls, although meditators have more broadband power

We compared the FC for the meditators relative to their matched control, during the eyes-open non-meditative state (EO1). As shown previously, meditators had more power at frequencies above ∼20 Hz in all electrodes relative to controls, even during non-meditative states (see Figure 2 in our previous study (Biswas et al., 2026)), which was also reflected in the PSD comparison plots in Figure 2a (frequencies for which the power was significantly different are shown in gray (p<0.05; uncorrected) or black (p<0.01; uncorrected)). Surprisingly, unlike power, meditators did not show a significant difference in FC (Figure 2b) compared to controls. There appeared to be a slight reduction in FC for meditators compared with controls, but this difference was not significant for most frequency bins. The scalp maps of FC within the same frequency window showed little difference between the two groups (Figure 2d). Both groups showed a similar decline in FC with increasing inter-electrode distance from the occipital seed electrodes (Figure 2e). We found similar results during the eyes-closed state and other non-meditative protocols as well (not shown).

**Figure 2:**
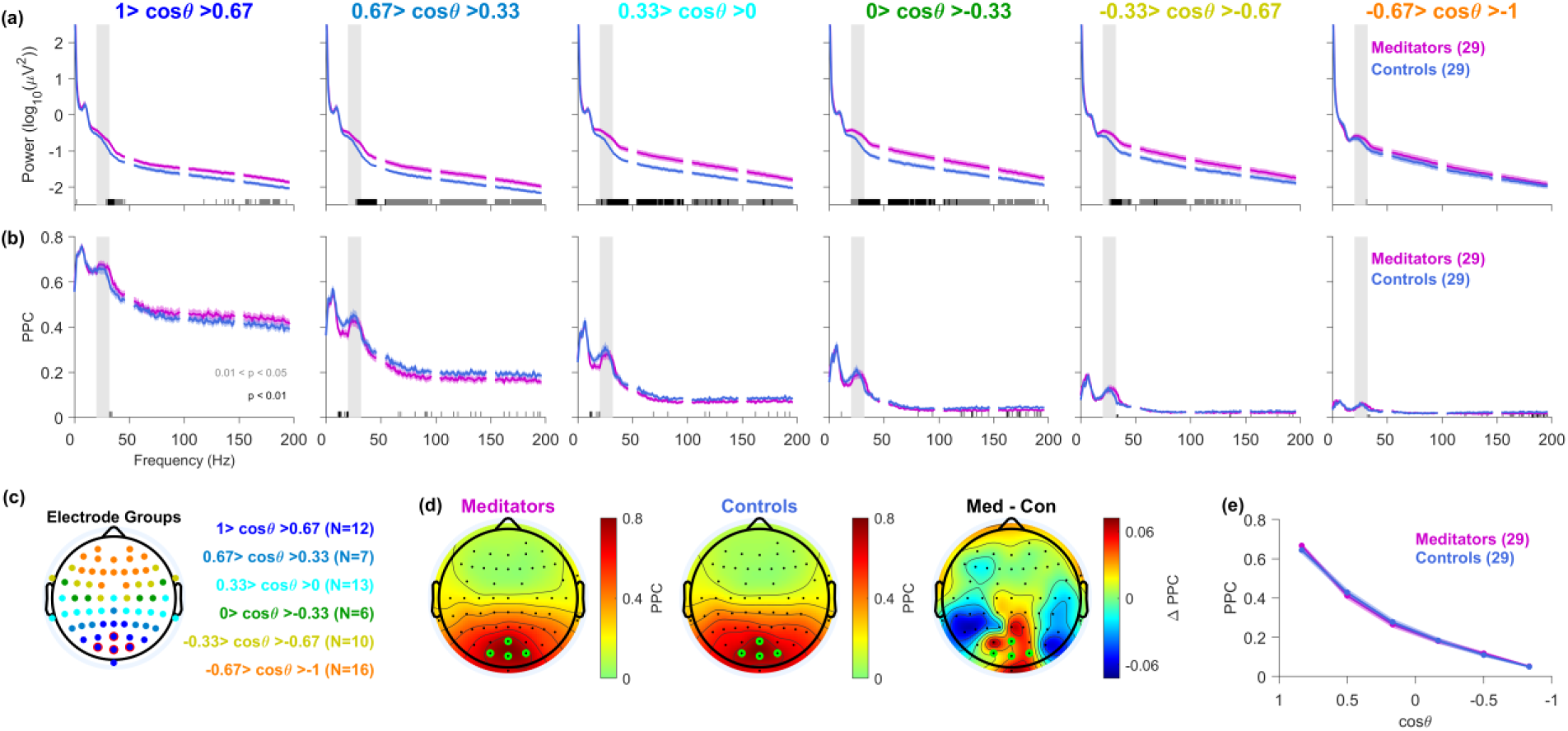
FC does not differ between meditators and controls during eyes open non-meditative state (EO1): **(a)** The PSDs of the six electrode groups averaged over the four seed electrodes and averaged across 29 meditators (magenta) and their matched controls (blue) for the EO1 protocol. The points around the line noise (47-53 Hz) and its harmonics are not shown. Solid traces represent the mean, and the shaded region around them indicates the standard error of the mean (SEM) across participants. The black and gray bars on the X-axis represent the significance of the difference in the mean of two groups (gray: p<0.05 and black: p<0.01, paired t-test). **(b)** The corresponding average FC relative to the occipital seed group – Oz, O1, POz, and O2 (highlighted in red in (c)) for the meditators (magenta) and their matched controls (blue). **(c)** The scalp layouts showing the six electrode groups defined based on the inter-electrode angular distance from the corresponding seed electrode, highlighted in red (for details, refer to‘Visualizing average FC’ under the’Methods’ section). The electrode groups are color-coded in terms of the cosine of their angular distance (cosθ) from the seed electrodes. **(d)** Average scalp FC for the meditators, controls, and the difference (meditators - controls), respectively, averaged over the frequency window (20-32) Hz, highlighted by the vertical shaded gray region in (a) and (b), and averaged over the four seed electrodes (highlighted in green). **(e)** Average FC for the meditators (magenta) and their matched controls (blue) plotted against inter-electrode distance from the seed electrodes. The FC value for each electrode group is obtained by averaging the PPCs in (b) over the frequency window of interest, 20-32 Hz.

As observed for the representative participant in Figure 1, the FC for both groups also showed a prominent‘bump’ in the beta range between 20-32 Hz (Figure 2b). In our previous studies, we used full-screen gratings that induce gamma oscillations in two distinct frequency bands, termed slow (20-34 Hz) and fast (35 – 70 Hz) gamma (Murty et al., 2018), and showed that slow gamma was stronger in meditators than controls, both before, after, and during meditation (Biswas et al., 2026). Even though the bump in the FC plots in Figure 2b falls within a similar frequency range, it is distinct from slow gamma as it is observed in the absence of a visual stimulus or meditation. We therefore refer to this stimulus-free activity as “beta” to distinguish it from the stimulus-induced slow-gamma that will be discussed later. Remarkably, this beta bump was prominent in FC plots even though it was not very prominent in the PSD plots (Figure 2a), suggesting that even during the resting state, there was enhanced beta FC throughout the cortex with respect to occipital electrodes. A similar bump (both in PSD and FC) was observed in the alpha range as well.

### Meditation is associated with a broadband suppression in FC

We next examined the effect of meditation (M1 and M2) on FC. Surprisingly, we found that meditation reduced FC over a broad frequency window (15-200 Hz and beyond), showing the opposite effect compared to PSD (Figure 3). The difference in FC across the two groups was most prominently observed in intermediate distance electrodes (2^nd^ and 3^rd^ electrode groups, 0 < cosθ < 0.67) relative to the occipital seed electrodes.

**Figure 3:**
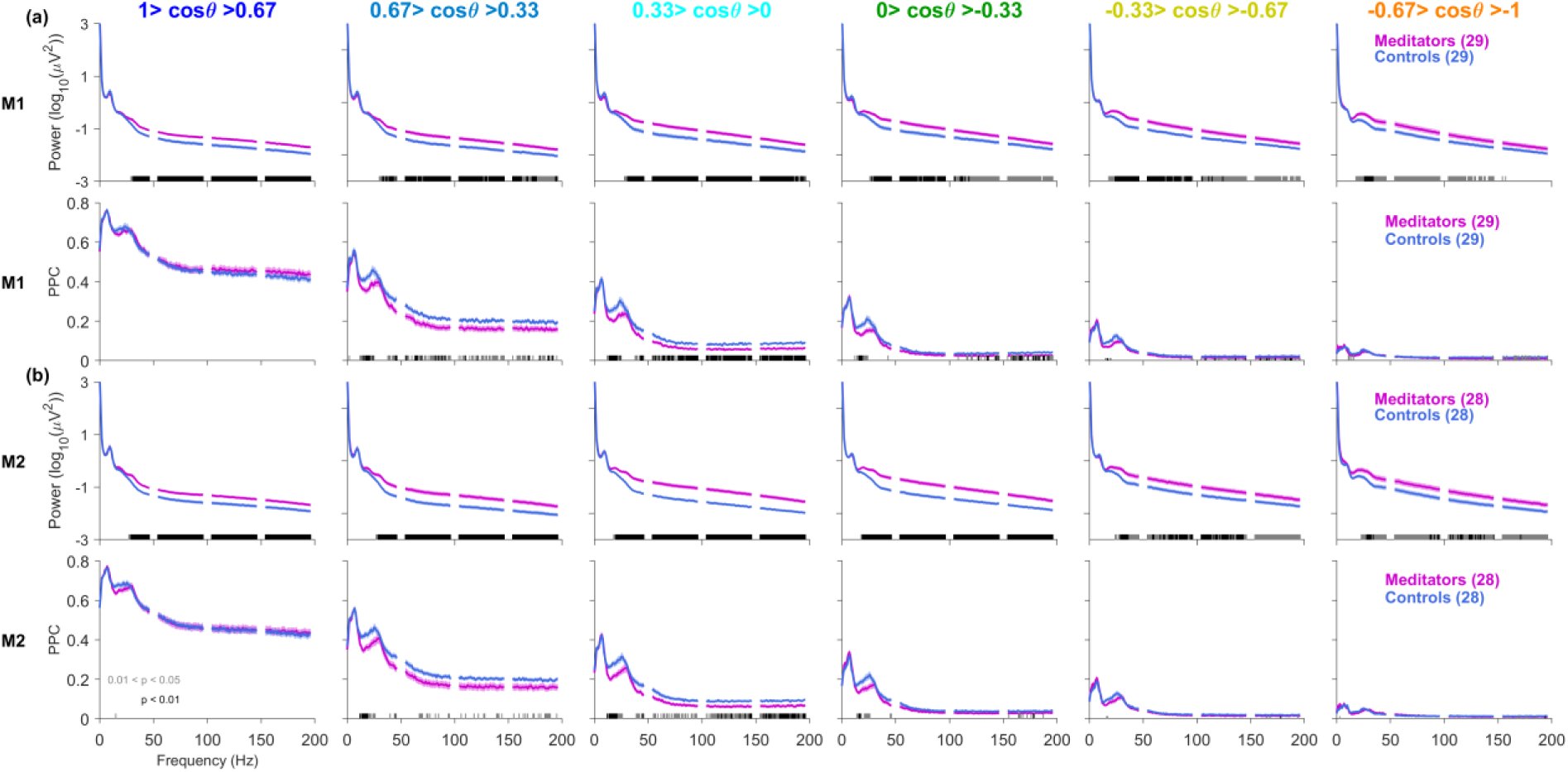
Meditation reduces functional connectivity (FC) across a broad frequency range: (a) top row: The PSD of the six electrode groups averaged across 29 meditators (magenta) and their matched controls (blue) during the M1 protocol. The seed electrodes (not shown here) remain the same as in Figure 2. Solid traces represent the mean, and the shaded region around them indicates the standard error of the mean (SEM) across participants. The black and gray bars on the X-axis represent the significance of the difference in the mean of two groups (gray: p<0.05 and black: p<0.01, paired t-test). **(a) bottom row:** The corresponding average FC relative to the occipital seed group of 29 meditators (magenta) and their paired controls (blue). **(b)** The same as (a) during the M2 protocol (n=28).

To further investigate the effect of meditation on FC, we examined the FC of meditators and controls separately during M1 (meditative state) relative to EO1. Interestingly, we found a significant drop in connectivity for the meditators but not for the controls, as shown in Figure 4a. To better visualize the effect of meditation across brain regions, we plotted scalp maps of FC averaged over the beta range (20-32 Hz) during M1 relative to the EO1 segment (Figure 4b; this is obtained by subtracting the FC topoplots for the two conditions). We found that the suppression in FC during meditation was much stronger in the temporal regions (Figure4b), which are located at interelectrode distances (0.67 to-0.67 in cosθ space) from the occipital seed electrodes (Figure4c top), although the reduction was significant for all six interelectrode distances (Figure 4c top; the p-values for the six electrode groups were 0.003, 3.954×10^-4^, 1.487×10^-4^, 0.005, 1.452×10^-4^ and 0.004 respectively; paired t-test; n=28). Similar results were observed during the baseline periods of M2 (not shown). For controls, the differences were not significant except for the first group that showed a slight increase (Figure 4c bottom; the p-values for the six electrode groups were 0.010, 0.532, 0.646, 0.701, 0.874, and 0.232, respectively; paired t-test; n=28).

**Figure 4:**
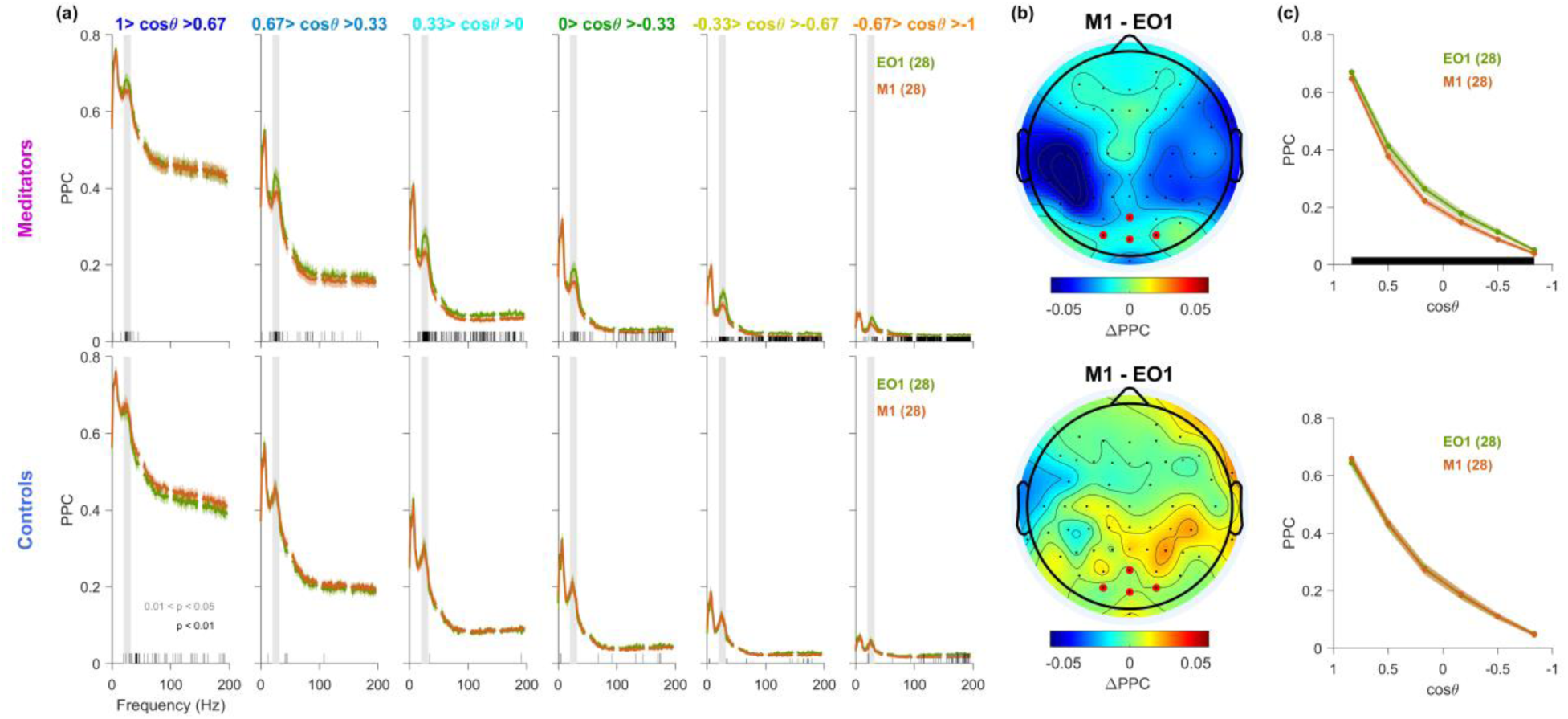
Meditation is associated with reduced beta FC: **(a)** The average FC of six electrode groups relative to the occipital seed group averaged across 28 meditators (top) and their matched controls (bottom) during M1 (orange) and EO1 (green). Solid traces represent the mean, and the shaded region around them indicates the standard error of the mean (SEM) across participants. The black and gray bars on the X-axis represent the significance of the difference in the mean of two groups (gray: p < 0.05 and black: p < 0.01, paired t-test). **(b)** Average difference in scalp FC between M1 and EO1 for meditators (top) and their matched controls (bottom) relative to occipital seed electrodes (highlighted in red) averaged over the frequency window (20-32 Hz), which is highlighted in the gray shaded region in (a). **(c)** Average FC for the meditators (top) and controls (bottom) during M1 (brown) and EO1 (green) plotted against inter-electrode distance from the seed electrodes. The FC value for each electrode group is obtained by averaging the PPCs in (a) over the frequency window of interest, 20-32 Hz. The black bars along X-axis represent the significance of group difference after Bonferroni correction (i.e., significant if p-values are less than 0.05/6, paired t-test; uncorrected p-values are reported in the text).

Further, we examined FC relative to the frontal seed electrodes: F1, F2, AF3, AF4, AFz, and Fz. We observed a similar broadband suppression in FC with meditation across the brain. Meditators exhibited a near-global reduction in FC during M1 compared to EO1 across a broad frequency range (15-200 Hz). Controls did not show any significant change in FC relative to frontal seed electrodes (Figure S1).

### Visual stimulation suppresses FC in both meditators and controls

Next, we studied the effect of visual stimulation on FC in both groups. We plotted FC for both stimulus (0.25 to 1.25 seconds) and baseline periods (-1 to 0 seconds, where 0 indicates stimulus onset) of the G1 protocol when visual gratings were shown to the participants (between 0 to 1.25 seconds). We saw that visual gratings, which enhance narrow-band gamma power (20-70 Hz) (Murty and Ray, 2022), led to reduced FC in both meditators and controls, as shown in Figure 5 (G1, meditators in top row and controls in bottom row, n=29; results are shown only up to 80 Hz to highlight the suppression in the beta range; the FC traces were highly overlapping and not significantly different above this range). This suppression was particularly significant in the beta range (20-32 Hz; highlighted in gray shadow in figures 5a and b). We found that this reduction in FC was more prominent in the temporal and parietal regions for both groups (Figure 5b). Both groups showed similar FC falloff with inter-electrode distance from the occipital seed electrodes (Figure 5c), with significant differences at all inter-electrode distances (p-values for meditators: 0.003, 0.001, 2.183 ×10^-4^, 4.960 ×10^-5^, 8.142 ×10^-5^, and 0.001, n=29, paired t-test; p-values for controls: 0.0034, 5.915 ×10^-4^, 1.468 ×10^-4^, 0.002, 0.002, and 0.026, n=29, paired t-test). We observed similar results for G2 and M2 protocols (not shown). Thus, unlike meditation, stimulus presentation only suppresses FC in a narrow band.

**Figure 5:**
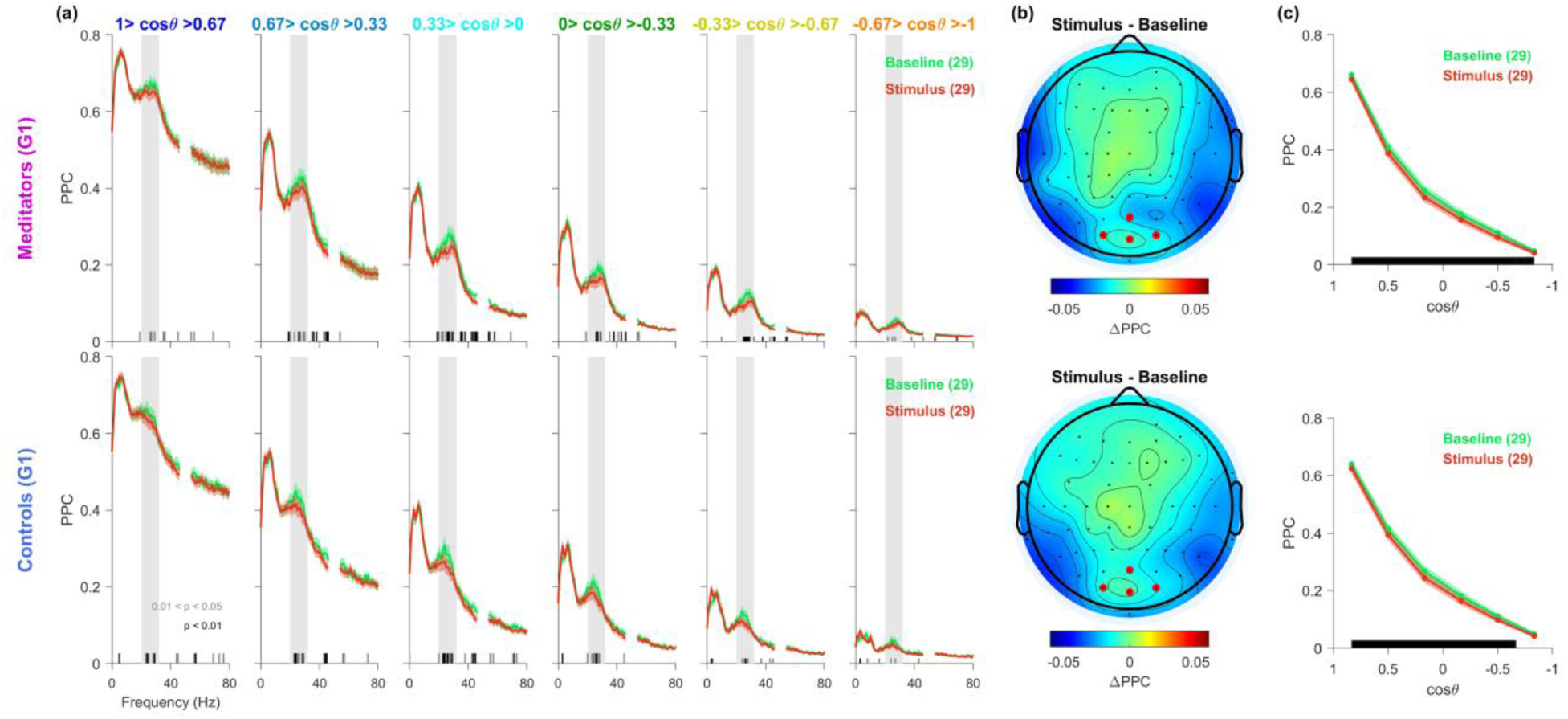
Presentation of the visual stimulus (grating) reduces FC in both meditators and controls: **(a)** Average FC of six electrode groups relative to the occipital seed group of meditators (top) and controls (bottom) during stimulus (red) and the baseline period of the trial (green) (n=29). Solid traces represent the mean, and the shaded region around them indicates the standard error of the mean (SEM) across participants. The black and gray bars on the X-axis represent the significance of the difference in the mean of two groups (gray: p < 0.05 and black: p < 0.01, paired t-test). **(b)** Average scalp FC for the meditators (top) and their matched controls (bottom) relative to occipital seed electrodes (highlighted in red) during stimulus compared to baseline averaged over the frequency window (20-32 Hz), which is shown in the gray shaded region in (a). **(c)** Average FC for the meditators (top) and their matched controls (bottom) during stimulus (red) and baseline (green) with inter-electrode distance from the seed electrodes. The FC value for each electrode group is obtained by averaging the PPCs in (a) over the frequency window of interest, 20-32 Hz. The black bars along X-axis represent the significance of group difference after Bonferroni correction.

### Effect of aging and gender on FC

We have previously found that healthy aging weakens stimulus-induced gamma power (Murty et al., 2020) and reduces FC in alpha and slow gamma ranges (Kumar and Ray, 2023). However, we studied FC only during the stimulus period to analyze stimulus-induced gamma oscillations and therefore did not analyze FC for the baseline period and across all frequencies. We therefore used the same dataset as the previous study (Murty et al., 2020, 2021; Murty and Ray, 2022; Kumar and Ray, 2023), in which participants performed a visual fixation task, where full-screen grating stimuli were presented for 800 ms with an inter-stimulus interval of 700 ms after a brief fixation of 1000 ms in each trial (Murty et al., 2020; Kumar and Ray, 2023).

Although there was no significant difference in baseline PSDs between middle-aged (50-64 years) and elderly participants (**≥** 65 years) (Figure 6a), we observed that the elderly participants, on average, showed reduced FC in both alpha (8-12 Hz) and high-beta (28-36 Hz) frequency ranges (Figure 6b). Similar to what we observed in meditators and controls, baseline FC showed a prominent bump in 20-36 Hz in both groups; however, the group difference emerged later on the‘falling edge’ of‘beta bump’ (28-36 Hz) (Figure 6b). Visual stimulus did not alter this effect, as previously reported (Kumar and Ray, 2023). The scalp maps show that the difference in FC in the same frequency window was larger in temporal regions (Figure 6d). The elderly group showed a suppressed FC falloff with inter-electrode distance from the occipital seed electrodes compared to the middle-aged group (Figure 6e; p values for the six electrode groups: 0.007, 0.009, 0.005, 5.856×10^-4^, 0.002 and 0.008; unpaired t-test).

**Figure 6:**
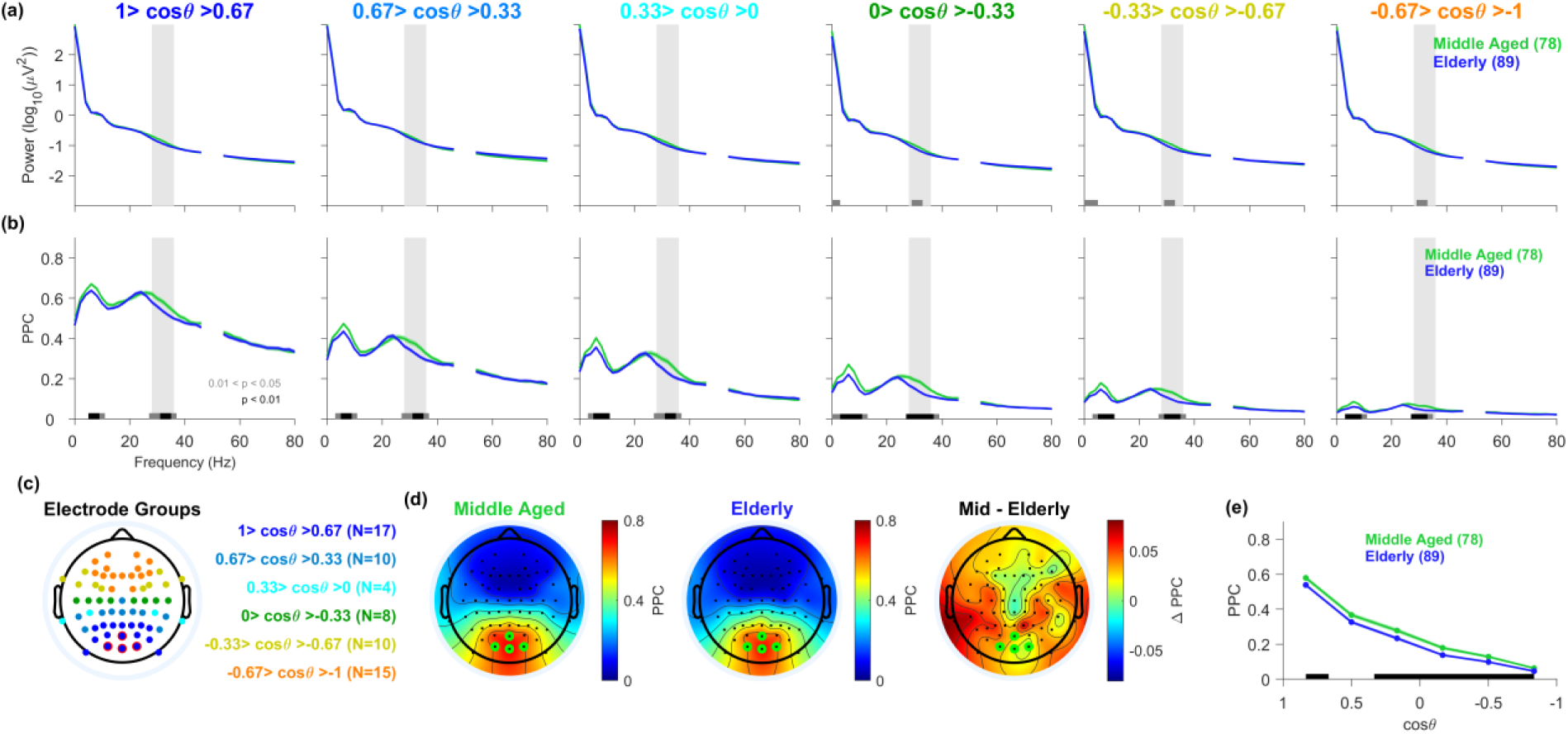
The effect of aging: **(a)** The baseline PSDs of the six electrode groups averaged across elderly (age > 64 years) (green, n=78) and middle-aged participants (age between 50 and 64 years) (blue, n=89) from the TLSA dataset. The points around the line noise (47-53 Hz) are not shown. Solid traces represent the mean, and the shaded region around them indicates the standard error of the mean (SEM) across participants. The black and gray bars on the X-axis represent the significance of the difference in the mean of two groups (gray: p < 0.05 and black: p < 0.01, paired t-test). **(b)** The corresponding average FC relative to the occipital seed group – Oz, O1, POz, and O2 (highlighted in red in (c)) of elderly participants (green) and middle-aged participants (blue). **(c)** Same as Figure 2c. **(d)** Average scalp FC for the elderly, middle-aged participants, and the difference (mid – elderly), respectively, averaged over the frequency window (28-36) Hz, highlighted by the vertical shaded gray region in (a) and (b), and averaged over the four seed electrodes (highlighted in green). **(e)** Average FC for the elderly (green) and middle-aged participants (blue) plotted against inter-electrode distance from the seed electrodes. The FC value for each electrode group is obtained by averaging the PPCs in (b) over the frequency window of interest, 28-36 Hz. The black bars along X-axis represent the significance of group difference after Bonferroni correction.

Interestingly, we also observed a gender difference in the FC in the same dataset. It has been known that males have weaker stimulus-induced gamma oscillations than females (Murty et al., 2020). In addition, we found that males have lower baseline FC than females for frequencies up to 60 Hz, but more prominently in the‘falling edge’ of the high-beta range (28-40 Hz) and mainly for relatively short inter-electrode distances relative to the occipital group (Figure S2; p values for the six electrode groups: 0.016, 0.003, 0.035, 0.024, 0.186, and 0.913; unpaired t-test). Similar results were obtained when the analysis was done during the stimulus period.

### Average referencing alters FC profiles

The analyses discussed so far were carried out on unipolar referenced signals. Previous studies have advocated using average referencing to avoid issues related to the choice of the reference electrode and spurious FC due to volume condition and bias from a single reference electrode (Srinivasan et al., 2007; Lehmann et al., 2012; Ríos-Herrera et al., 2019). We therefore reperformed the above analyses after average re-referencing of the signals. While differences in power between meditators and controls persisted, the broadband suppression in FC was no longer observed (Figure 7). Moreover, with increasing interelectrode distance, PPC no longer decreased monotonically, but paradoxically increased beyond a certain distance (equivalent to cos θ <-0.33; Figure 7). To explain this discrepancy, we plotted polar histograms of phase differences between seed and other electrodes, after pooling phasors across trials, seed electrodes, electrodes within a group, frequency range between 20-32 Hz, and across participants (meditators or controls) for both unipolar and average-referenced signals (Figure 8). For unipolar referencing, the distribution was skewed toward 0°, hence the mean phasor direction was near 0° for all six electrode groups in both meditators and controls. The phase distribution broadened with increasing inter-electrode distance, such that the length of the mean phasor (i.e., phase locking value or PLV), monotonically decreased with inter-electrode distance in both meditators and controls, as expected (Figure 8a). But for the average re-referenced signals, the distribution of phase differences flipped from the mean phasor direction of ∼0° to ∼180° with increasing interelectrode distance for both meditators and controls (Figure 8b). The PLV initially decreased with increasing interelectrode distance, vanished at cos θ ∼ 0 (at an angular distance ∼ 90°), and increased in the opposite direction with increasing distance from the seed electrode. Although in both referencing schemes, meditators, on average, had smaller PLV than controls, this group difference was significant only for unipolar referenced signals. We also tested this for each participant individually (not shown) and the same pattern was observed for most participants. We discuss the reasons for this paradoxical behaviour later.

**Figure 7:**
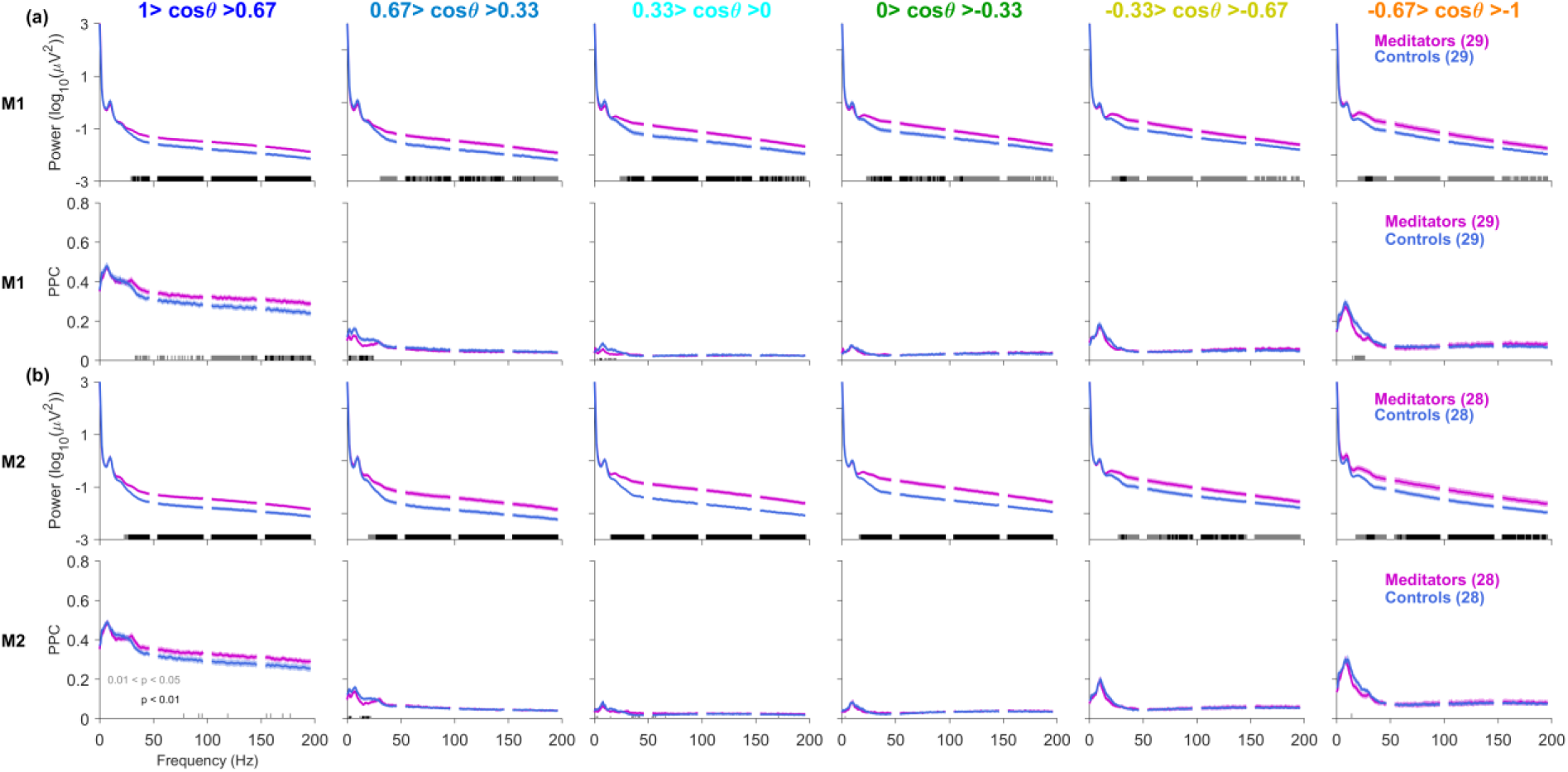
Average re-referencing results in no significant group difference between meditators and controls: (a) top row: The average-referenced PSD of the six electrode groups averaged across 29 meditators (magenta) and their matched controls (blue) during the M1 protocol. The occipital seed electrodes (not shown here) remain the same as in Figure 2. Solid traces represent the mean, and the shaded region around them indicates the standard error of the mean (SEM) across participants. The black and gray bars on the X-axis represent the significance of the difference in the mean of two groups (gray: p < 0.05 and black: p < 0.01, paired t-test). **(a) bottom row:** The corresponding average FC relative to the occipital seed group of 29 meditators (magenta) and their paired controls (blue). **(b)** The same as (a) during the M2 protocol (n=28).

**Figure 8:**
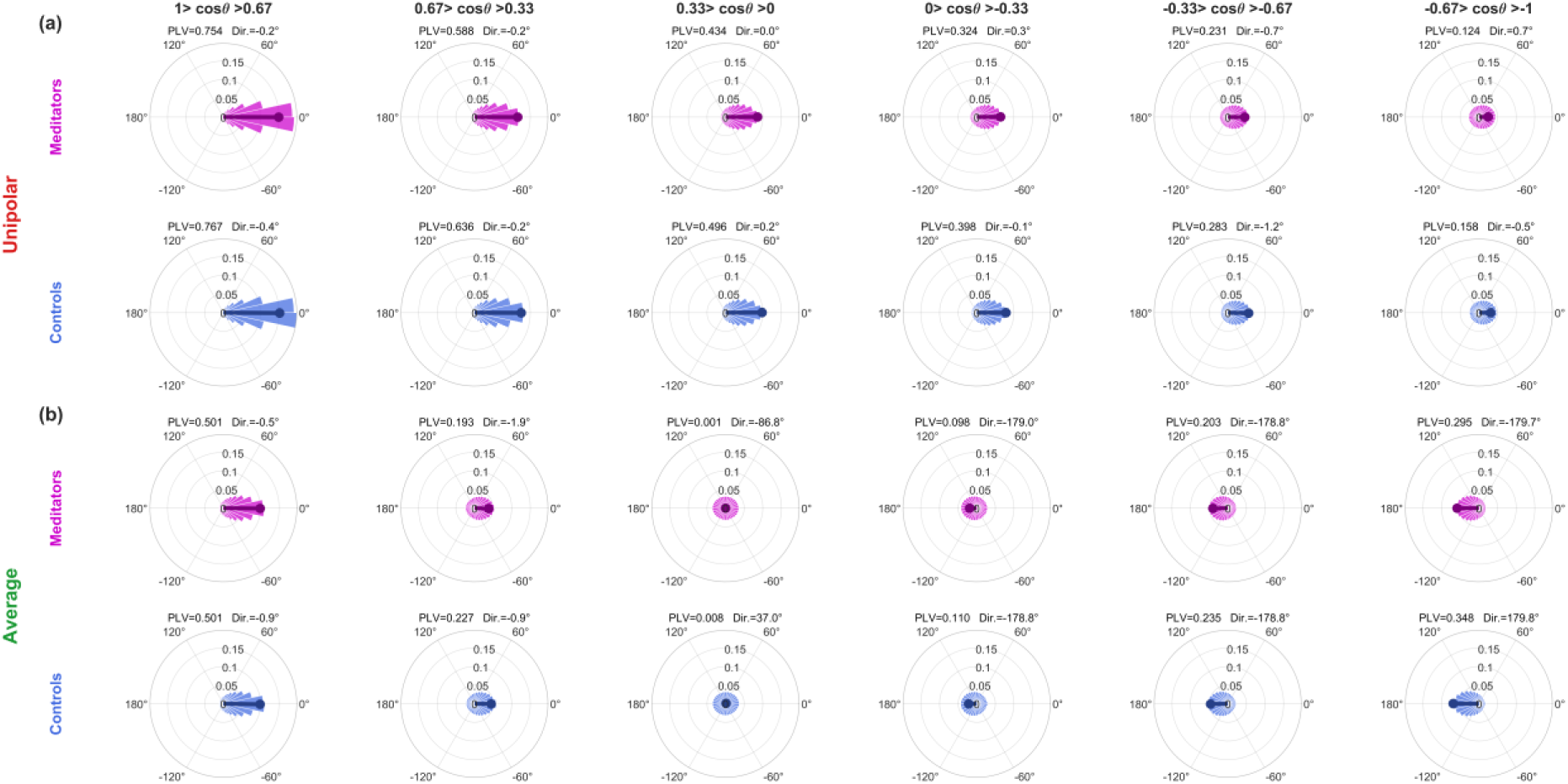
Average re-referencing alters the distribution of phase differences: **(a)** Unipolar referenced signals: Polar histograms of phase differences between occipital seed electrodes (same as Figure 2c) and other electrodes pooled across the six electrode groups, four seed electrodes, the frequency range 20-32 Hz, and participants – 30 meditators (magenta, top row) and 30 controls (blue, bottom row) during the M1 protocol. The occipital seed electrodes (not shown here) and the corresponding six electrode groups remain the same as in Figure 2. The average phasor is shown in dark shade. Its length, i.e., PLV and direction are mentioned just above each rose plot. The maximum global radius across the rose plots is 0.1976. The PLV vector is scaled down by the same factor. **(b)** The same as (a) for average referenced signals.

Other FC metrics have been suggested that are less prone to volume conduction, such as imaginary coherence (Nolte et al., 2004), wPLI (Vinck et al., 2011), and lagged coherence (Pascual-Marqui, 2007). In all these metrics, some variant of the imaginary part of coherency is used and the synchronous component (vectors pointing towards zero degrees) is ignored. However, as shown in Figure 8, the mean directions of vectors always remained at either 0°or 180° irrespective of the inter-electrode distance. Therefore, imaginary coherence (icoh) during M1 and M2 was close to zero and largely invariant with interelectrode distance (Figure S3) and did not yield a significant group difference between meditators and controls at any frequency. Similar results were obtained for wPLI and lagged coherence (not shown).

## Discussion

We examined changes in EEG FC with meditation, and compared meditation-induced changes with changes due to visual stimulus, healthy aging, and gender. Across protocols and seed electrode groups, FC spectra consistently exhibited peaks in the alpha (8–12 Hz) and beta (∼20–32 Hz) bands, suggesting that large-scale neuronal coupling is intrinsically stronger in these frequency ranges. Although meditators showed elevated broadband power (15–200 Hz) even at rest and further increase in power during meditation (Biswas et al., 2026), FC did not differ between meditators and controls during non-meditative states. However, meditation induced a broadband suppression of FC, particularly in temporal-occipital pairs. In contrast, stimulus, aging, and gender effects were more “narrowband” in nature: visual stimulation reduced beta FC globally, aging reduced alpha and beta FC, and females exhibited elevated high-frequency FC (notably 28–40 Hz) than males. Importantly, common-average referencing severely altered the FC profiles, shifting the average phase difference from ∼0**°** to ∼180**°** as the interelectrode distance increased.

### Meditation-induced broadband‘state’ suppression of FC

Overall reduced FC during meditation (M1 and M2) across a broad frequency range (15-200 Hz) may possibly reflect reduced self-referential processing and less automatic mind-wandering (Berkovich-Ohana et al., 2014), linked with the deactivation of the default mode network (DMN) (Gusnard et al., 2001; Christoff et al., 2009). Similarly, another study reported reduced lagged coherence in theta, alpha, and beta bands across multiple meditation traditions (Lehmann et al., 2012), aligning with our findings of decreased FC, although our effects extended over a broader frequency range and depended on the use of PPC computed on data that was not average referenced. In contrast, another study reported increased long-range gamma synchrony (25–42 Hz) during compassion meditation (Lutz et al., 2004). These discrepancies may reflect differences in meditation style (compassion versus Rajyoga) and in the FC metric (wavelet-derived phase synchrony versus PPC), suggesting that “meditation” is not a unitary neural state.

The beta-bump in FC is worth noting. Beta oscillations are often linked to sensorimotor functions such as postures and movements, and top-down inhibitory control (Quivira-Lopesino et al., 2025). Since our reference electrode was also in the sensory-motor region, one possibility is that the default beta FC bump is due to this choice of referencing. However, the absence of a corresponding beta peak in PSD argues against a purely reference-driven artifact. Average referencing eliminated the beta FC peak but also simultaneously distorting the FC spectrum.

Irrespective of the origins of the beta FC, the suppression of broadband FC with meditation (and absence of movements) in Figure 4 suggests reduced neural synchrony spanning a wide frequency range. Importantly, although reduced beta FC has been associated with impaired cognition in clinical populations (Shah-Basak et al., 2019), such an interpretation is unlikely here, given the well-documented cognitive benefits of meditation (Sumantry and Stewart, 2021). Instead, it is more likely to reflect the release of top-down inhibitory control in an effortless meditative state and can be viewed as a breakdown of long-range neuronal coupling, allowing distant networks to operate more independently, as previously interpreted by others as well (Jensen and Mazaheri, 2010).

It is worth noting that muscular activity can potentially contaminate the EEG activity above 20 Hz (Whitham et al., 2007). But EMG-driven changes would produce correlated changes in power and FC. Instead, we observed increased power but decreased FC during meditation, arguing against this explanation. This dissociation supports the view that power and FC capture distinct features of ongoing neural activity (Siems et al., 2016): power reflects local excitability and E–I balance (Bartos et al., 2007; Buzsáki and Wang, 2012), whereas FC reflects large-scale coordination (Lachaux et al., 1999).

The deviations in our results from other meditation studies reporting covariance of power and FC could also arise from the stark difference in meditation style (Transcendental vs Rajyoga). Also, it is important to note that power and FC changed in different directions with meditation and stimulus; they co-varied when comparing age and gender (e.g., middle-aged participants had more power as well as higher FC as compared to elderly participants; same for females versus males).

Microsaccades can artefactually induce broadband high-frequency activity and spurious FC (Yuval-Greenberg et al., 2008; Keren et al., 2010), but their contribution here is likely minimal: FC differences appeared during meditation (M1) but not in a comparable eyes-open baseline (EO1), and gaze was limited to 2.5° about the fixation spot. Additionally, although coherence decreases with increasing uncorrelated noise (Artieda et al., 2006), our extensive artifact rejection minimizes this concern.

### Stimulus-induced reduction in FC

Visual stimuli induce “narrowband” gamma oscillations in the visual cortex (Murty and Ray, 2022) and reduce alpha-band synchronization, especially over occipital and parietal regions (Pfurtscheller and Lopes da Silva, 1999), potentially due to disruption of the resting-state-synchronized alpha network (Barry et al., 2009). We observed that a grating stimulus led to a slight reduction in FC in the beta band globally, particularly in temporoparietal-occipital pairs, in both meditators and controls (Figure 5). This is also supported by our previous finding that change in power and FC are negatively correlated in the slow-gamma band (Kumar and Ray, 2023). This might reflect the functional decoupling of visual input from the cortical regions, especially in the posterior areas. The fact that this effect was similar in meditators and controls suggests that it is largely stimulus-driven and independent of long-term meditation practice, likely reflecting intrinsic sensory processing dynamics rather than top-down cognitive control.

### The effect of healthy aging and gender

Healthy aging is associated with decreased alpha power (Scally et al., 2018; Murty et al., 2020) and increased broadband high-frequency power, with a flattening of PSD, often interpreted as increased neural noise (Vysata et al., 2012; Voytek et al., 2015; Aggarwal and Ray, 2023). Prior studies have reported reductions in FC across delta, theta, alpha, and gamma bands, alongside increases in beta FC (Moezzi et al., 2019).

In our data, unlike meditation, aging was associated with reduced narrowband alpha and high-beta (28-40 Hz) FC, particularly in posterior networks, even when power differences were minimal (Figure 6). These effects were present both during stimulus presentation and inter-stimulus intervals, suggesting that they reflect stable trait-like changes rather than task-specific modulation. Reduced FC may indicate diminished efficiency of large-scale integration and weakened top-down control. This interpretation is consistent with reports of reduced posterior alpha connectivity in aging (Perinelli et al., 2022) and broader frameworks describing age-related changes in network integration and segregation (Kavčič et al., 2023).

Both power and FC showed gender differences. Females showed higher global power in the 15-60 Hz range and increased centroparietal–occipital FC in the 28-40 Hz band. Similar findings have been reported previously: females exhibit higher global wPLI during sleep (Ujma et al., 2019) and greater coherence during verbal memory tasks (Volf and Razumnikova, 1999). These differences may reflect sex-dependent neurodevelopmental or anatomical organization (Volf and Razumnikova, 1999; McCarthy, 2016; Ujma et al., 2019). Importantly, we emphasize that these group-level electrophysiological differences should not be overinterpreted in terms of cognitive or behavioral superiority of either gender.

### Average referencing disrupts the phase relationship between signals

Referencing had a profound impact on FC estimates. With common-average referencing, group differences between meditators and controls were no longer significant (Figure 7), and overall FC magnitude decreased. Further, average referencing produced non-monotonic effects: the average phaser shortened initially, then lengthened, and the phase distribution shifted from 0° toward 180° with increasing interelectrode distance, effectively flipping phase relationships (Figure 8). Polar hostograms of the phase-difference distribution revealed clustering of phasors near 0° and 180° (Figure 8), explaining the remarkably reduced magnitudes of phase-lag-based metrics such as imaginary coherence (Figure S3), wPLI, and lagged coherence.

This phenomenon has been demonstrated previously (Shirhatti et al., 2016) in LFP signals. It arises because direct spectral estimators are themselves noisy and yield amplitude estimates that follow a Rayleigh distribution and power that follows an exponential distribution (Srinath and Ray, 2014)). Therefore, many of the phasors are smaller than the amplitude of the average reference, and subtraction of average reference signal inverts the phase by ∼180° (Shirhatti et al., 2016). Subtracting the global average from all signals can also eliminate meaningful shared signal components and thereby distort FC (Essl and Rappelsberger, 1998). Therefore, caution is warranted when applying average referencing in phase-based metric analyses.

One of the limitations of our analyses is the use of sensor space for FC estimation. It is believed that FC in the cortical source space is more reliable, less prone to volume conduction, and reflects true functional patterns underlying cognitive processes (Lehmann et al., 2012; Xie et al., 2022). However, it is a standard practice to use average referencing before doing source localization (as done, for example, in eLORETA). In addition, methods such as LORETA also employ a Laplacian operator, which also distorts the phase distribution (Shirhatti et al., 2016; the “CSD reference” used there is the same as a Laplacian operator where the average of the four neighboring electrodes is used as the reference signal). Thus, more systematic analysis is needed to test whether FC spectrum is distorted when analysis is done on the source space.

Our results, overall, suggest that the long-term practice of meditation modulates cortical neural activity, both power and FC, in a manner distinct from those of healthy aging and visual stimulation. Our findings can help identify functional patterns of healthy aging, network abnormalities underlying cognitive impairments, and inform the development of meditation-based therapeutic interventions.

## Supplementary Figures

**Figure S1:**
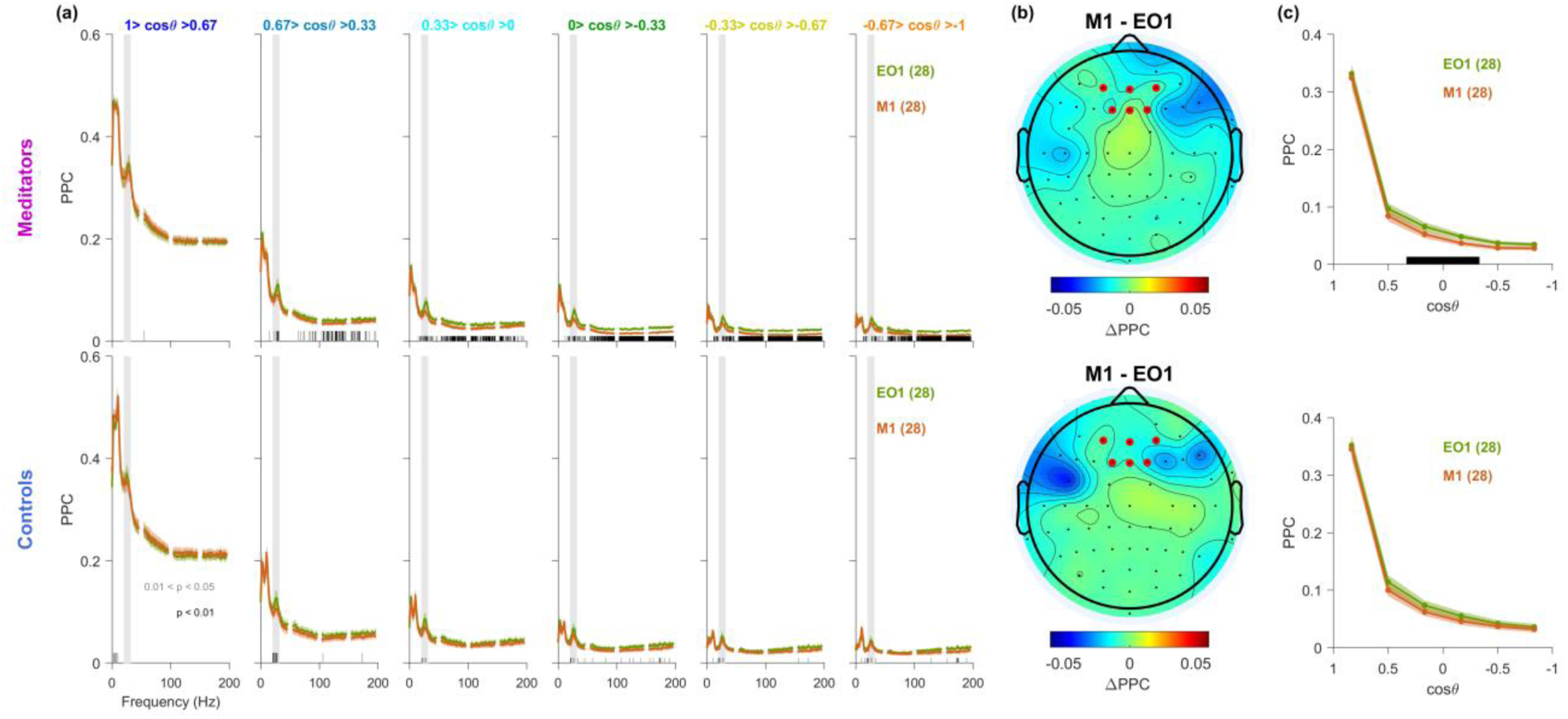
Frontal connectivity: Same as Figure 5, but when seed electrodes are chosen from frontal area (shown in red dots in c).

**Figure S2:**
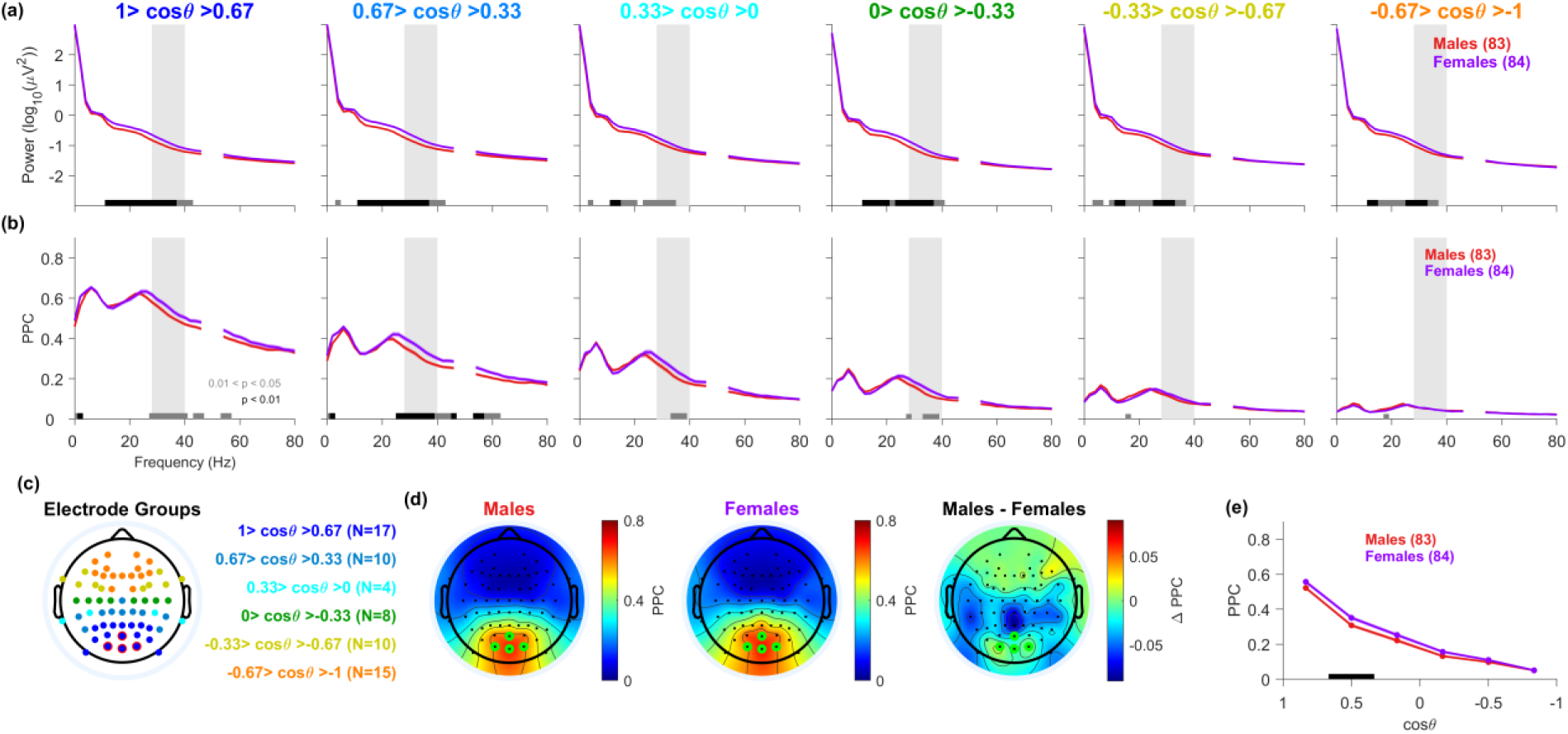
The gender difference in FC: Results are shown in the same format as Figure 6, but compared between males (83) and females (84).

**Figure S3:**
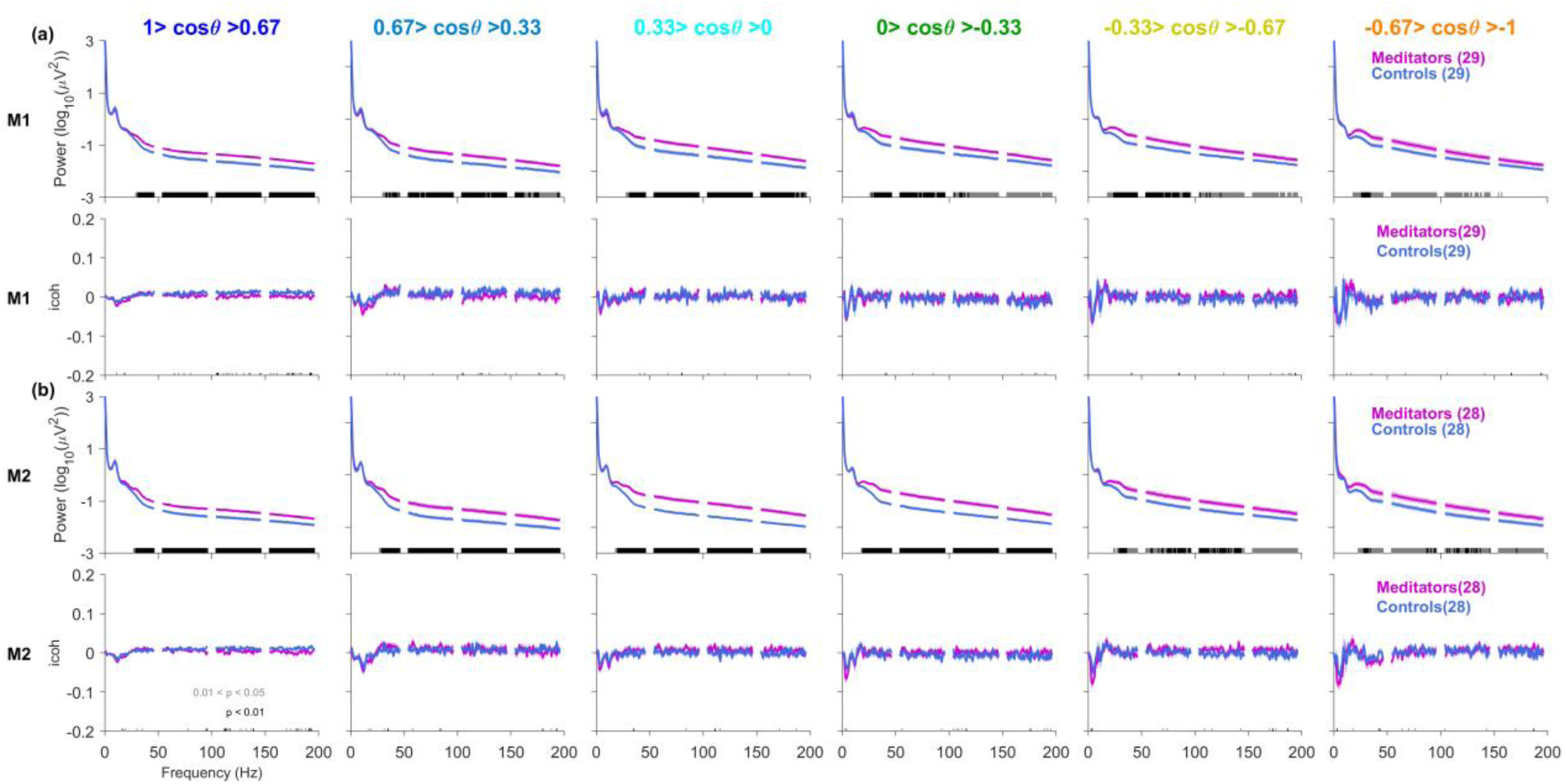
Imaginary coherence (iCoh) for meditators and controls shows no statistically significant difference: Same as Figure 3 but using iCoh instead of PPC.

## Notes

**Funding Disclosure** This work was supported by DBT/Wellcome Trust India Alliance (Senior fellowship IA/S/18/2/504003) and a grant from Pratiksha Trust to S.R., DST INSPIRE Fellowship to V.J., and Institute Gate and Axis Bank PhD fellowship to A.B.

### Competing Interest Statement

The authors have declared no competing interest.

## References

Aggarwal S, Ray S (2023) Slope of the power spectral density flattens at low frequencies (<150 Hz) with healthy aging but also steepens at higher frequency (>200 Hz) in human electroencephalogram. Cereb Cortex Commun 4:tgad011.

Artieda J, Alegre M, Valencia M (2006) Electric Fields of the Brain. The Neurophysics of EEG, second ed. Paul L. Nunez, Ramesh Srinivasan. Clin Neurophysiol - CLIN NEUROPHYSIOL 117:2109–2110.

Barry RJ, Clarke AR, Johnstone SJ, Brown CR (2009) EEG differences in children between eyes-closed and eyes-open resting conditions. Clin Neurophysiol 120:1806–1811.

Bartos M, Vida I, Jonas P (2007) Synaptic mechanisms of synchronized gamma oscillations in inhibitory interneuron networks. Nat Rev Neurosci 8:45–56.

Berkovich-Ohana A, Glicksohn J, Goldstein A (2014) Studying the default mode and its mindfulness-induced changes using EEG functional connectivity. Soc Cogn Affect Neurosci 9:1616–1624.

Biswas A, Aggarwal S, Sharma K, Ray S (2026) Simultaneous enhancement of stimulus-induced and stimulus-free gamma in open-eye meditators. Imaging Neurosci 4:IMAG.a.1145.

Bokil H, Andrews P, Kulkarni JE, Mehta S, Mitra PP (2010) Chronux: a platform for analyzing neural signals. J Neurosci Methods 192:146–151.

Braboszcz C, Cahn BR, Levy J, Fernandez M, Delorme A (2017) Increased Gamma Brainwave Amplitude Compared to Control in Three Different Meditation Traditions. PloS One 12:e0170647.

Buzsáki G, Wang X-J (2012) Mechanisms of Gamma Oscillations. Annu Rev Neurosci 35:203–225.

Cahn BR, Polich J (2006) Meditation states and traits: EEG, ERP, and neuroimaging studies. Psychol Bull 132:180–211.

Christoff K, Gordon AM, Smallwood J, Smith R, Schooler JW (2009) Experience sampling during fMRI reveals default network and executive system contributions to mind wandering. Proc Natl Acad Sci U S A 106:8719–8724.

Delorme A, Makeig S (2004) EEGLAB: an open-source toolbox for analysis of EEG dynamics. J Neurosci Methods 134:9–21.

Essl M, Rappelsberger P (1998) EEG coherence and reference signals: experimental results and mathematical explanations. Med Biol Eng Comput 36:399–406.

Ferrarelli F, Smith R, Dentico D, Riedner BA, Zennig C, Benca RM, Lutz A, Davidson RJ, Tononi G (2013) Experienced Mindfulness Meditators Exhibit Higher Parietal-Occipital EEG Gamma Activity during NREM Sleep. PLoS ONE 8:e73417.

Gusnard DA, Akbudak E, Shulman GL, Raichle ME (2001) Medial prefrontal cortex and self-referential mental activity: relation to a default mode of brain function. Proc Natl Acad Sci U S A 98:4259–4264.

Huang Y, Zhang J, Cui Y, Yang G, He L, Liu Q, Yin G (2017) How Different EEG References Influence Sensor Level Functional Connectivity Graphs. Front Neurosci 11:368.

Ivanovski B, Malhi GS (2007) The psychological and neurophysiological concomitants of mindfulness forms of meditation. Acta Neuropsychiatr 19:76–91.

Jensen O, Mazaheri A (2010) Shaping Functional Architecture by Oscillatory Alpha Activity: Gating by Inhibition. Front Hum Neurosci 4:186.

Kavčič A, Demšar J, Georgiev D, Bon J, Soltirovska-Šalamon A (2023) Age related changes and sex related differences of functional brain networks in childhood: A high-density EEG study. Clin Neurophysiol 150:216–226.

Kayser J, Tenke CE (2015) Issues and considerations for using the scalp surface Laplacian in EEG/ERP research: A tutorial review. Int J Psychophysiol Off J Int Organ Psychophysiol 97:189–209.

Keren AS, Yuval-Greenberg S, Deouell LY (2010) Saccadic spike potentials in gamma-band EEG: Characterization, detection and suppression. NeuroImage 49:2248–2263.

Kumar WS, Ray S (2023) Healthy ageing and cognitive impairment alter EEG functional connectivity in distinct frequency bands. Eur J Neurosci 58:3432–3449.

Lachaux JP, Rodriguez E, Martinerie J, Varela FJ (1999) Measuring phase synchrony in brain signals. Hum Brain Mapp 8:194–208.

Lehmann D, Faber PL, Tei S, Pascual-Marqui RD, Milz P, Kochi K (2012) Reduced functional connectivity between cortical sources in five meditation traditions detected with lagged coherence using EEG tomography. NeuroImage 60:1574–1586.

Lomas T, Ivtzan I, Fu CHY (2015) A systematic review of the neurophysiology of mindfulness on EEG oscillations. Neurosci Biobehav Rev 57:401–410.

Lutz A, Greischar LL, Rawlings NB, Ricard M, Davidson RJ (2004) Long-term meditators self-induce high-amplitude gamma synchrony during mental practice. Proc Natl Acad Sci U S A 101:16369–16373.

McCarthy MM (2016) Multifaceted origins of sex differences in the brain. Philos Trans R Soc B Biol Sci 371:20150106.

Miskovic V, Keil A (2015) Reliability of event-related EEG functional connectivity during visual entrainment: Magnitude squared coherence and phase synchrony estimates. Psychophysiology 52:81–89.

Moezzi B, Pratti LM, Hordacre B, Graetz L, Berryman C, Lavrencic LM, Ridding MC, Keage HAD, McDonnell MD, Goldsworthy MR (2019) Characterization of Young and Old Adult Brains: An EEG Functional Connectivity Analysis. Neuroscience 422:230–239.

Murty DV, Manikandan K, Kumar WS, Ramesh RG, Purokayastha S, Nagendra B, Ml A, Balakrishnan A, Javali M, Rao NP, Ray S (2021) Stimulus-induced gamma rhythms are weaker in human elderly with mild cognitive impairment and Alzheimer’s disease. eLife 10:e61666.

Murty DVPS, Manikandan K, Kumar WS, Ramesh RG, Purokayastha S, Javali M, Rao NP, Ray S (2020) Gamma oscillations weaken with age in healthy elderly in human EEG. NeuroImage 215:116826.

Murty DVPS, Ray S (2022) Stimulus-induced Robust Narrow-band Gamma Oscillations in Human EEG Using Cartesian Gratings. Bio-Protoc 12:e4379.

Murty DVPS, Shirhatti V, Ravishankar P, Ray S (2018) Large Visual Stimuli Induce Two Distinct Gamma Oscillations in Primate Visual Cortex. J Neurosci Off J Soc Neurosci 38:2730–2744.

Nair AK, Sasidharan A, John JP, Mehrotra S, Kutty BM (2017) Just a minute meditation: Rapid voluntary conscious state shifts in long term meditators. Conscious Cogn Int J 53:176–184.

Nolte G, Bai O, Wheaton L, Mari Z, Vorbach S, Hallett M (2004) Identifying true brain interaction from EEG data using the imaginary part of coherency. Clin Neurophysiol 115:2292–2307.

Oostenveld R, Fries P, Maris E, Schoffelen J-M (2011) FieldTrip: Open source software for advanced analysis of MEG, EEG, and invasive electrophysiological data. Comput Intell Neurosci 2011:156869.

Pascual-Marqui RD (2007) Coherence and phase synchronization: generalization to pairs of multivariate time series, and removal of zero-lag contributions. Available at: https://ui.adsabs.harvard.edu/abs/2007arXiv0706.1776P [Accessed April 21, 2026].

Perinelli A, Assecondi S, Tagliabue CF, Mazza V (2022) Power shift and connectivity changes in healthy aging during resting-state EEG. NeuroImage 256:119247.

Pfurtscheller G, Lopes da Silva FH (1999) Event-related EEG/MEG synchronization and desynchronization: basic principles. Clin Neurophysiol 110:1842–1857.

Quivira-Lopesino A, Sevilla-García M, Cuesta P, Pusil S, Bruña R, Fiedler P, Cebolla AM, Cheron G, Funke M, Maestu F (2025) Changes of EEG beta band power and functional connectivity during spaceflight: a retrospective study. Sci Rep 15:13399.

Ríos-Herrera WA, Olguín-Rodríguez PV, Arzate-Mena JD, Corsi-Cabrera M, Escalona J, Marín-García A, Ramos-Loyo J, Rivera AL, Rivera-López D, Zapata-Berruecos JF, Müller MF (2019) The Influence of EEG References on the Analysis of Spatio-Temporal Interrelation Patterns. Front Neurosci 13 Available at: https://www.frontiersin.org/journals/neuroscience/articles/10.3389/fnins.2019.00941/full [Accessed January 4, 2026].

Scally B, Burke MR, Bunce D, Delvenne J-F (2018) Resting-state EEG power and connectivity are associated with alpha peak frequency slowing in healthy aging. Neurobiol Aging 71:149–155.

Shah-Basak PP, Dunkley BT, Ye AX, Wong S, da Costa L, Pang EW (2019) Altered beta-band functional connectivity may be related to‘performance slowing’ in good outcome aneurysmal subarachnoid patients. Neurosci Lett 699:64–70.

Sharma K, Achermann P, Panwar B, Sahoo S, Pascual-Marqui R, Faber P, Ramakrishnan AG (2023) High Theta–Low Alpha Modulation of Brain Electric Activity During Eyes-Open Brahma Kumaris Rajyoga Meditation. Mindfulness.

Shen Y-Q, Zhou H-X, Chen X, Castellanos FX, Yan C-G (2020) Meditation effect in changing functional integrations across large-scale brain networks: Preliminary evidence from a meta-analysis of seed-based functional connectivity. J Pac Rim Psychol 14:e10.

Shirhatti V, Borthakur A, Ray S (2016) Effect of Reference Scheme on Power and Phase of the Local Field Potential. Neural Comput 28:882–913.

Siems M, Pape A-A, Hipp JF, Siegel M (2016) Measuring the cortical correlation structure of spontaneous oscillatory activity with EEG and MEG. NeuroImage 129:345–355.

Sifis M, Vourkas M, Tsirka V, Karakonstantaki E, Kanatsouli K, Stam C (2009) The Influence of Ageing on Complex Brain Networks: A Graph Theoretical Analysis. Hum Brain Mapp 30:200–208.

Srinath R, Ray S (2014) Effect of amplitude correlations on coherence in the local field potential. J Neurophysiol 112:741–751.

Srinivasan R, Winter WR, Ding J, Nunez PL (2007) EEG and MEG coherence: measures of functional connectivity at distinct spatial scales of neocortical dynamics. J Neurosci Methods 166:41.

Sumantry D, Stewart KE (2021) Meditation, Mindfulness, and Attention: a Meta-analysis. Mindfulness 12:1332–1349.

Sumner RL, McMillan RL, Shaw AD, Singh KD, Sundram F, Muthukumaraswamy SD (2018) Peak visual gamma frequency is modified across the healthy menstrual cycle. Hum Brain Mapp 39:3187–3202.

Tang Y-Y, Ma Y, Wang J, Fan Y, Feng S, Lu Q, Yu Q, Sui D, Rothbart MK, Fan M, Posner MI (2007) Short-term meditation training improves attention and self-regulation. Proc Natl Acad Sci 104:17152–17156.

Thomson D (1982) Spectrum Estimation and Harmonic Analysis. Proc IEEE 70:1055–1096.

Ujma PP, Konrad BN, Simor P, Gombos F, Körmendi J, Steiger A, Dresler M, Bódizs R (2019). Sleep EEG functional connectivity varies with age and sex, but not general intelligence. Neurobiol Aging 78:87–97.

Vinck M, Oostenveld R, van Wingerden M, Battaglia F, Pennartz CMA (2011) An improved index of phase-synchronization for electrophysiological data in the presence of volume-conduction, noise and sample-size bias. NeuroImage 55:1548–1565.

Vinck M, van Wingerden M, Womelsdorf T, Fries P, Pennartz CMA (2010) The pairwise phase consistency: a bias-free measure of rhythmic neuronal synchronization. NeuroImage 51:112–122.

Volf NV, Razumnikova OM (1999) Sex differences in EEG coherence during a verbal memory task in normal adults. Int J Psychophysiol Off J Int Organ Psychophysiol 34:113–122.

Voytek B, Kramer MA, Case J, Lepage KQ, Tempesta ZR, Knight RT, Gazzaley A (2015) Age-Related Changes in 1/f Neural Electrophysiological Noise. J Neurosci 35:13257–13265.

Vysata O, Kukal J, Prochazka A, Pazdera L, Valis M (2012) Age-Related Changes in the Energy and Spectral Composition of EEG. Neurophysiology 44:63–67.

Whitham EM, Pope KJ, Fitzgibbon SP, Lewis T, Clark CR, Loveless S, Broberg M, Wallace A, DeLosAngeles D, Lillie P, Hardy A, Fronsko R, Pulbrook A, Willoughby JO (2007) Scalp electrical recording during paralysis: quantitative evidence that EEG frequencies above 20 Hz are contaminated by EMG. Clin Neurophysiol Off J Int Fed Clin Neurophysiol 118:1877–1888.

Xie W, Toll RT, Nelson CA (2022) EEG functional connectivity analysis in the source space. Dev Cogn Neurosci 56:101119.

Yuval-Greenberg S, Tomer O, Keren AS, Nelken I, Deouell LY (2008) Transient Induced Gamma-Band Response in EEG as a Manifestation of Miniature Saccades. Neuron 58:429–441.

